# Neural Crest-Derived Endocardial Cells Required for Heart Valve Development

**DOI:** 10.64898/2026.11.03.763802

**Authors:** Alexander F. Merriman, Mauro W. Costa, Angelo Pelonero, Yu Huang, Tomohiro Nishino, Feiya Li, Shyam Lal Jinagal, Adrienne Lam, Abigail Brand, Victoria Yu, Yan Hu, Feng Gao, Irfan S. Kathiriya, Shafkat Anwar, Michael Bunker, Sanjeev S. Ranade, Casey A. Gifford, Benoit G. Bruneau, Chun Jimmie Ye, Deepak Srivastava

## Abstract

Abnormalities in cardiac valves, which maintain unidirectional blood flow through the heart, affect 2–3% of people worldwide and are the most common form of structural heart disease^1^.Yet the cell types that make up the valve, and how they communicate to direct development and homeostasis, remain poorly understood. Here, we identify a rare, essential population of highly secretory valve endocardial cells in mouse and human that derive from migrating neural crest cells. Ablation of these cells in mice caused perinatal lethality with thickened leaflets, stenosis, and regurgitation, mirroring human valve disease. This subset of endocardial cells uniquely expresses the secreted signaling protein parathyroid hormone-like hormone (*Pthlh*). Global disruption of Pthlh caused valve stenosis and lethality and neural crest-specific deletion produced similar valve disease, with impaired maturation of valvular interstitial cells and disrupted bone morphogenetic protein signaling. These findings identify a previously unrecognized cell type and signaling pathway essential for valve morphogenesis, with implications for the pathogenesis and treatment of human valve defects.

## Introduction

Disruption of normal cardiac valvulogenesis during embryonic development can cause congenital valvulopathies, which account for 20–30% of all congenital heart defects (CHD), including those with the highest morbidity and mortality rates^2,3^. Furthermore, adult valve abnormalities represent the third most common form of heart disease and often are due to abnormal formation or maintenance of valves. Development of the valvular precursors—the endocardial cushions—is one of the first events to occur during cardiogenesis to ensure unidirectional flow of blood. During murine development at embryonic day (E) 9.5, mesodermally-derived endocardial cells undergo an endocardial-to-mesenchymal transition (EMT) event at both the inflow and outflow regions of the developing heart in response to reciprocal signaling between specialized myocardium adjacent to valve endocardium. From E9.5 to E12.5, the valvular mesenchyme expands through the proliferation of transformed endocardial cells, in addition to contributions from the epicardium and cardiac neural crest mesenchyme. While many of the molecular regulators of initial cardiac valve emergence have been described, little is understood about post-EMT valvulogenesis. This later stage of valvulogenesis is of particular interest, as defects during this stage are more likely to give rise to human CHDs, whereas more severe defects earlier in development may result in embryonic lethality.

Recent studies have demonstrated the critical importance of the endocardium in the regulation of valvulogenesis and associated valvular defects in humans^4^. Moreover, there appears to be endocardial heterogeneity in postnatal valves^5^, although the developmental origin has been thought to be uniformly from early cardiac mesoderm in contrast to the developmental heterogeneity of valve mesenchyme. In this study, we utilized single-cell transcriptomic analysis of post-EMT embryonic mouse heart development to discover a previously unrecognized sub-population of endocardial cells unexpectedly derived from migratory neural crest cells. This population of highly secretory cells was essential for normal valvulogenesis, and disruption of secreted parathyroid hormone-like hormone (*Pthlh*) signaling in the neural crest lineage caused valve stenosis and regurgitation similar to human disease.

## Results

### *Pthlh* Expression Marks a Rare Valvular Endocardial Subpopulation

To characterize the cellular heterogeneity during post-EMT valvulogenesis, we performed single cell RNA sequencing (scRNA-seq) on mouse hearts at E12.5, the time point at which the endocardium, epicardium, and neural crest lineages have each contributed to the valvular mesenchyme^6^ (Fig. 1a). A total of 13,926 cells were annotated using expression of well-known marker genes (Fig 1b, Extended Data Fig. 1a - c). Valvular endocardium (*Nfatc1*^7^ and *Klf2*^8^), valvular mesenchyme (*Sox9*^9^ and *Postn*^10^), and specialized valvular myocardium (*Rspo3*^11^ and *Tbx2*^12^) were re-clustered into 7 distinct sub-types revealing surprising heterogeneity within populations (Extended Data Fig. 1d,e). To investigate ligand– receptor interactions between these cell types during post-EMT valvulogenesis, we used the computational pipeline CellChat, performing a global cell-cell communication pattern analysis based on the expression of paracrine ligands from sender cells and their cognate receptors in receiver cells^13^. Among the most striking predictions was a valvular endocardial cell (VEC) subtype as a sender of Hedgehog^14^ and Wnt^15^ signaling, along with the parathyroid hormone-like hormone (*Pthlh*) signaling pathway. While the importance of Hedgehog and Wnt signaling in valvulogenesis has been reported, *Pthlh*, a peptide that can function as a hormone or paracrine factor, has not previously been investigated in valvulogenesis or cardiogenesis (Fig. 1c). Network centrality analysis of the Pthlh pathway identified a putative interaction between the VEC-derived parathyroid hormone-like hormone (PTHLH) ligand and parathyroid hormone 1 receptor (*Pth1r*)-expressing valvular mesenchyme and valvular myocardium (Fig. 1d, Extended Data Fig. 1f). *Pthlh* is an important regulator of endochondral bone formation^16^—a developmental process that has considerable overlap with valvulogenesis based on transcriptomic analysis^17^.

**Figure 1.**
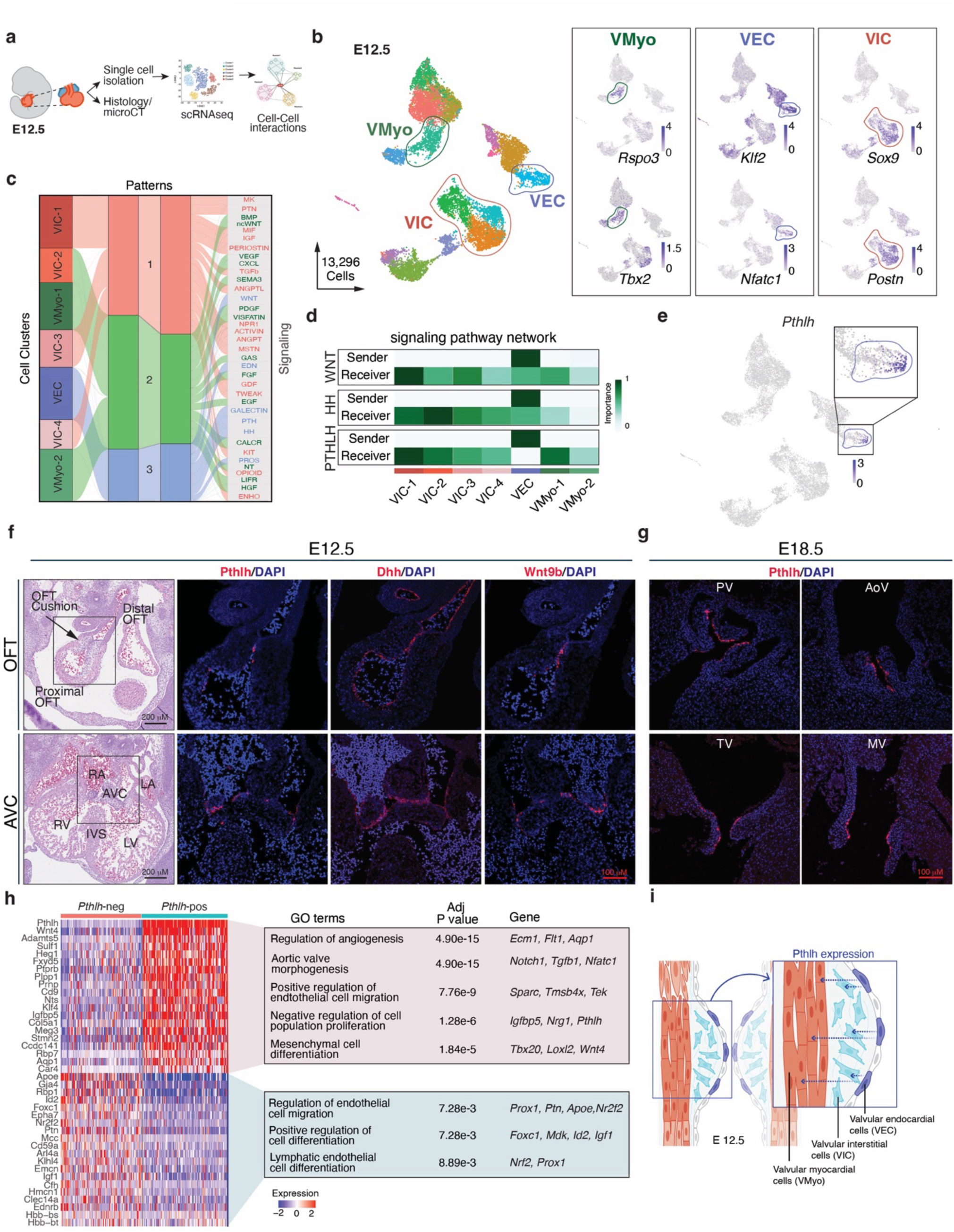
*Pthlh* marks a novel, highly secretory subset of valvular endocardium that is spatiotemporally-specific to post-EMT valvulogenesis. **a**, Schematic of experimental approach to micro-dissect and dissociate E12.5 mouse hearts and perform downstream cell-cell interaction analysis. **b,** (left) Annotated Uniform manifold approximation and projection (UMAP) plot from scRNA-seq of whole mouse hearts at E12.5 (n = 2 WT), colored by cluster identity, with the valvular myocardium (VMyo, green circle), valvular endocardial cells (VEC, blue circle), and valvular interstitial cells (VIC, red circle) highlighted. (right) UMAP plots for previously characterized marker gene expression for *Rspo3, Tbx2, Klf2, Nfatc1, Sox9,* and *Postn*. **c**, River plot of predicted secretory patterns in the developing E12.5 valves as predicted by CellChat. **d**, Network centrality analysis heatmaps predicting sending and receiving cell types in the developing valve for the *Wnt*, *HH*, and *PTHLH* signaling pathways. **e**, UMAP plot of *Pthlh* expression with the valvular endocardial cell cluster highlighted (blue circle) and a zoomed in view in the right upper corner. **f**, Hematoxylin and eosin staining (left column) and fluorescent *in situ* hybridization for *Pthlh, Dhh*, and *Wnt9b* from serial adjacent sections of the outflow tract (OFT) (top row) and atrioventricular canal (AVC) (bottom row) at E12.5. **g**, Fluorescent *in situ* hybridization for *Pthlh* from sections of the pulmonary valve (PV), aortic valve (AoV) (top row), and tricuspid valve (TV) and mitral valve (MV) (bottom row) at E18.5. **h**, Heatmap plot of differentially expressed genes between *Pthlh*^+^/*Cdh5*^+^ and *Pthlh*^-^/*Cdh5*^+^ cells (adjusted *P* value < 0.05 & Log2FC >0.75). Top GO terms enriched in up- and down-regulated genes with representative genes with corresponding adjusted *P* values for each GO term represented in adjacent boxes. **i,** Cartoon schematic demonstrating *Pthlh*-secreting endocardium to *Pth1r*-expressing valvular myocardium and mesenchyme at E12.5.

*Pthlh* transcripts at E12.5 were restricted to a small subset of *Nfatc1^+^/Klf2^+^* VECs (Fig. 1e), representing less than 1% of cells in the developing heart. Spatially, *Pthlh*^+^ expression was found in a limited number of endocardial cells of both the atrioventricular and outflow tract cushions most distal to the valvular myocardium and nearest the opposing cushion (Fig. 1f). *Pthlh^+^* cells and endocardial cells co-expressed *Wnt9b* and *Dhh* in scRNAseq data (Extended Data Fig. 1g), and this was validated by RNAscope on tissue sections (Fig. 1f). At E18.5, *Pthlh*^+^ cells remained confined to the coaptation surface of the valve leaflets in a punctate pattern across all four valves (Fig. 1g). Thus, in the developing heart, *Pthlh* expression marks a relatively small sub-population of spatially-restricted valvular endocardium.

To determine when *Pthlh*^+^ cells first emerge during cardiogenesis, we assembled an atlas of scRNAseq datasets at several time points during cardiogenesis from De Soysa et al.^18^, from Ranade et al.^19^, and from the current study (Extended Data Fig 2a - c). We found a robust increase in both the percentage of *Pthlh^+^* cells and the level of *Pthlh* expression between E10.5 and E11.5, temporally coinciding with the end of EMT and the progression into post-EMT valvulogenesis^6^ (Extended Data Fig. 2d). Genes expressed in E12.5 *Pthlh*^+^ cells displayed a high degree of concordance with marker genes for coaptation of VECs previously identified postnatally at post-natal day 7 (P7) and P30, suggesting that *Pthlh*^+^ cells persist into postnatal remodeling and adult life.

To define the transcriptional signature of this rare endocardial cell population compared to other endocardial cells, we performed differential gene expression analysis between *Pthlh*^+^/*Cdh5*^+^ and *Pthlh*^−^/*Cdh5*^+^ endocardium at E12.5 (Fig. 1h). Gene ontology (GO) analysis of the *Pthlh*^+^ cell transcriptional signature was enriched for genes related to aortic valve development, angiogenesis regulation, endothelial cell migration, and mesenchymal cell differentiation. Notably, relative to *Pthlh*^−^ endocardial cells, *Pthlh*^+^ cells had lower expression of important endocardial genes such as *Emcn*^26^ and *Ednrb*^20^, and genes related to lymphatic endothelial cell differentiation (e.g., *Prox1*), another postnatal valvular endocardial cell type that has been described^21^. Moreover, we observed an enrichment in the expression of numerous secreted developmental signaling ligands in *Pthlh*^+^/*Cdh5*^+^ vs. *Pthlh*^−^/*Cdh5*^+^ cells, including *Wnt4, Wnt9b*, *Wnt11, Dhh*, *Nppc*, several retinoic acid signaling regulators, and proteins involved in cell-cell junction maintenance (Fig. 1e; Extended Data Figs. 1c, 2e). Spatial transcriptomics analysis of adult mouse hearts showed that both *Pthlh* and its receptor Pth1r were present in mature aortic valves, with *Pthlh* expression still restricted to a subset of endocardial layers of the mature valves (Extended Data Fig. 2f). This unique transcriptional signature and expression profile raises interesting questions related to the function of this rare, highly secretory subpopulation during development (Fig. 1i).

To investigate the presence of *PTHLH*^+^ endocardial cells in human hearts, we analyzed published spatial data from embryonic hearts^22^ and found that PTHLH^+^ cells are also detected in the valve forming regions of 9 PCW fetal hearts, displaying a high degree of co-localization with CDH5 valve endocardial cells (Fig. 2a,b). Furthermore, single cell transcriptomic data from 9 human fetal heart samples collected between 90-120 days after fertilization revealed a subset of human *PTHLH^+^/CDH5^+^*endocardial cells (Fig. 2c) ^4,23^, together indicating the presence of a similar rare cell type in human valvulogenesis. To further probe possible role of PTHLH in disease, we performed scRNA-seq on a 20-week gestation human fetal heart with severe aortic valve stenosis/atresia and associated hypoplastic left heart syndrome (HLHS). All four valves were dissected, enzymatically and mechanically dissociated, and processed independently to preserve valve-specific identities (Extended Data Fig. 3a). Sequencing and downstream analysis yielded a 36,416-cell Seurat object comprising 22 clusters representing canonical valvular populations (Extended Data Fig. 3b,c). Despite overall transcriptional similarities across valves, the unique *PTHLH*^+^/CDH5^+^ endocardial subset was identifiable (Extended Data Fig. 3d), consistent with our prior findings in the human fetal heart cell atlas. Notably, the critically stenosed aortic valve showed marked depletion of *PTHLH*^+^ cells amongst CDH5^+^ endocardial cells relative to the other healthy valves (Extended Data Fig. 3e). These diseased *PTHLH*^+^ aortic valve cells had a unique transcriptional signature, including reduced natriuretic peptide precursor C (*NPPC*) expression relative to the *PTHLH*^+^ cells in the non-diseased pulmonary valve (Extended Data Fig. 3f,g). Degenerative aortic valve stenosis with calcification in adults is associated with a 90% reduction in *NPPC* gene expression levels within the aortic valve, raising the possibility that *PTHLH*^+^ cells may also play a role in adult aortic valve pathology^46^, warranting future analysis of additional fetal and adult valves.

**Figure 2.**
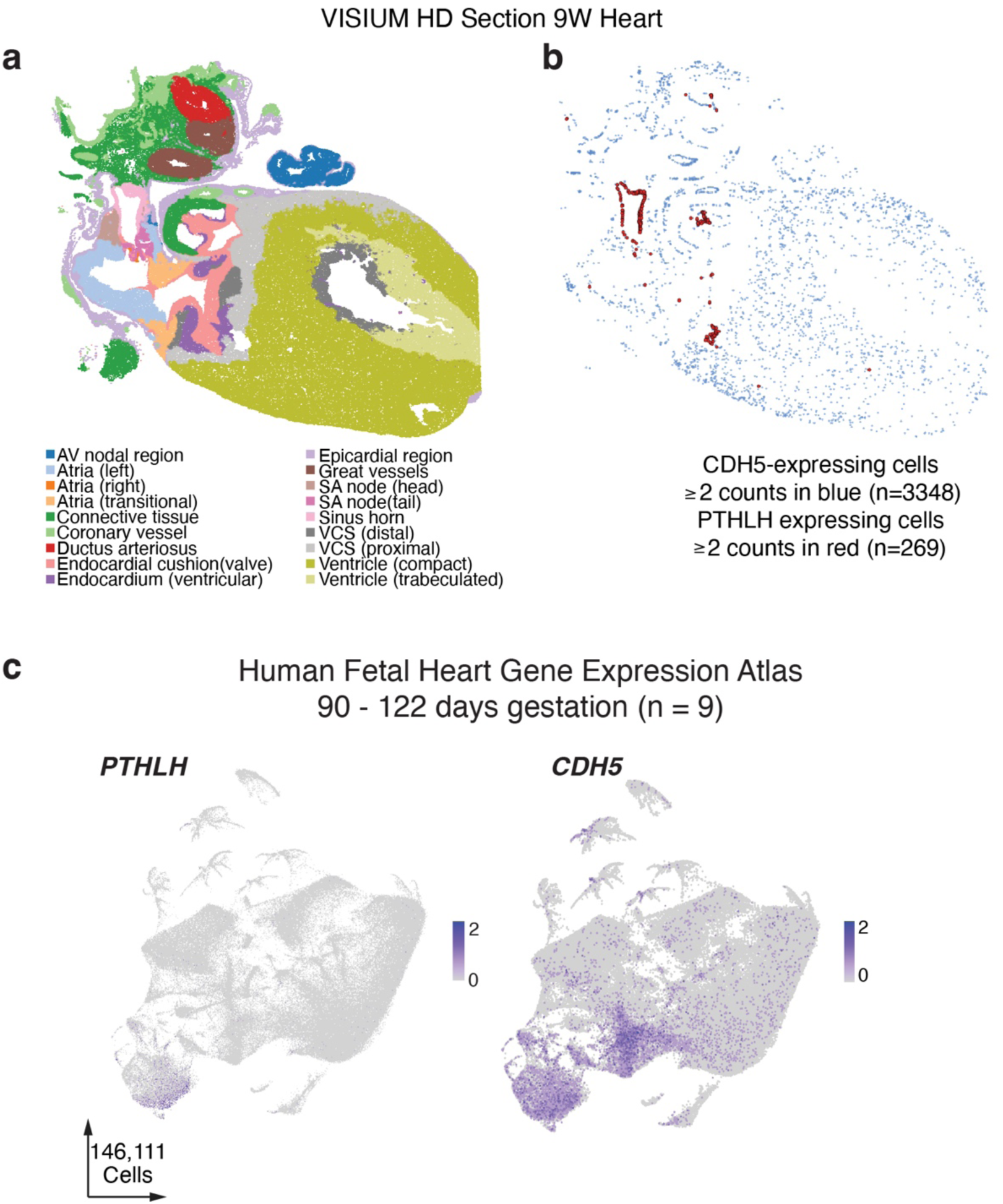
PTHLH-expressing cells in the human fetal heart. Visium HD spatial transcriptomics of a 9 post-conception-week (PCW) human heart (Cranley et al., 2026)^22^. **a**, tissue niches of the section (CellCharter, n = 136,685 cells). **b**, CDH5-expressing cells (blue, n = 3,348) and PTHLH-expressing cells (red, n = 269; >= 2 UMI). PTHLH-expressing cells are concentrated in the endocardial cushion/valve niche and the sinus horn. **c,** UMAP plot of PTHLH and CDH5 expression from 9 human fetal heart samples ranging from 90-122 days gestation, reprocessed from Cao et al.^23^.

### *Pthlh*^+^ Cells are Essential to Post-EMT Valvulogenesis

To test the requirement of *Pthlh*^+^ cells during valvulogenesis, we used a tamoxifen-inducible *PthlhCreER* mouse line^24^ intercrossed with a mouse conditionally expressing diphtheria toxin A (RosaDTA)^25^ to selectively kill *Pthlh*-expressing cells (Fig. 3a). We induced ablation just prior to the emergence of *Pthlh^+^* cells at E10.5, and continued daily tamoxifen injections until E14.5. Within hours of birth, initial litters (n = 3 litters) displayed perinatal cyanosis and lethality in one hundred percent of the ablated *PthlhCreER*^+^:*RosaDTA*^+^ pups (n = 8), whereas control littermates appeared healthy; this was similar to the perinatal lethality observed in whole-organism *Pthlh* knockout^26^. Fetal echocardiography at E18.5 revealed that *PthlhCreER*^+^:*RosaDTA*^+^ mice had turbulent flow across the aortic and pulmonary valves, indicative of congenital valvular stenosis (Fig. 3b). We additionally identified atrioventricular valvular regurgitation and occasional flow of blood from the left ventricle to the right ventricle just below the aortic valve, indicative of a sub-aortic ventricular septal defect (Extended Data Fig. 4a-c). This type of septal defect often occurs in humans in conjunction with aortic valve abnormalities. Histologic analysis revealed that *Pthlh*^+^ cell ablation caused valvular dysplasia characterized by thickened, immature- appearing leaflets, with the aortic and pulmonary valves most severely impacted. The penetrance of pulmonary or aortic valve dysplasia was 80% or 73.3%, respectively, concordant with the functional phenotypes observed via echocardiography (Fig. 3c, Extended Data Table 1). To validate our histologic findings and interrogate the phenotype in three-dimensions, we performed microCT on E18.5 mouse hearts^27^ and transformed images into three-dimensional models (Fig. 3d). These three-dimensional models recapitulated the phenotypic features identified via histologic analysis. Thus, loss of *Pthlh*^+^ cells resulted in perinatal cyanosis and death secondary to valvular dysplasia similar to that observed in human CHDs.

**Figure 3.**
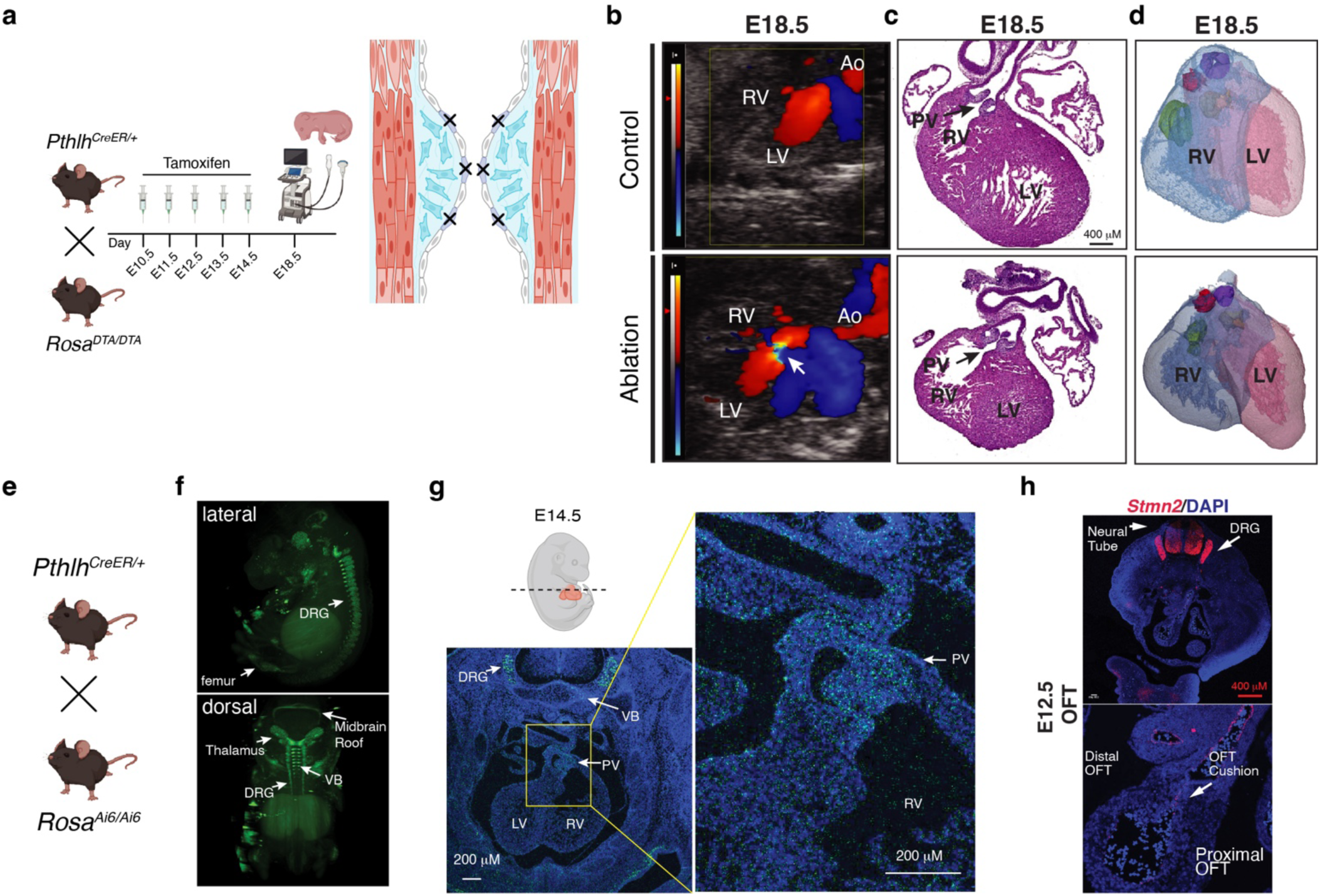
*Pthlh* valvular endocardial cells are essential for post-EMT valvulogenesis and derived from migratory neural crest. **a**, Strategy for inducible ablation of *Pthlh*^+^ cells. Rosa^DTA/DTA^ females were mated with PthlhCreER^+^; RosaDTA/^+^ males and were injected daily with tamoxifen from E10.5– 14.5, embryos were screened via fetal echocardiography at E18.5, and dissection and subsequent histologic analysis was performed comparing PthlhCreER/^+^;RosaDTA/DTA (Ablation) hearts against control littermates. **b**, Fetal echocardiogram from control (top) and ablation (bottom) E18.5 hearts. **c**, Hematoxylin and eosin staining of E18.5 control (top) and ablation (bottom) hearts, with a dysplastic pulmonary valve (PV) shown (Black arrow). **d**, Three-dimensional reconstruction of control (top) and ablation (bottom) E18.5 hearts from microCT data. Aortic valve (red), Pulmonary valve (Purple), Mitral valve (yellow), tricuspid valve (green). **e**, Mating strategy for *Pthlh*^+^ cell lineage tracing. PthlhCreER^+^ males were mated with Ai6 females; pregnant females were injected daily from E10.5–E14.5. **f**, Lightsheet micrograph of a cleared E14.5 embryo in lateral (top) and dorsal (bottom) views displaying endogenous ZsGreen fluorescence secondary to PthlhCreER–mediated recombination. **g**, Section *in situ* hybridization for ZsGreen (green) from an E14.5 embryo littermate in an approximate axial plane (black dashed lines in cartoon). **h**, Fluorescent *in situ* hybridization for *Stmn2* expression (red) in the OFT at 4x (top) and 20x (bottom) magnification at E12.5 in a wild-type embryo.

### Pthlh^+^ Cells are Neural Crest-Derived

Given the novelty of this rare subpopulation and its requirement in mice, we investigated the developmental origin and fate of *Pthlh*^+^ cells. We crossed the *Pthlh*^CreER^ mouse line to a *Rosa26^Ai6/Ai6^*fluorescent (ZsGreen) reporter line^28^, using the same TAM injection scheme as for the ablation model (Fig. 3e). Using lightsheet microscopy of cleared E14.5 embryos, we detected endogenous *ZsGreen* expression not only in the anticipated mesodermal structures (e.g., extremity long bones, vertebral bodies), but also unexpectedly in a number of neural-crest derived structures, including dorsal root ganglia (DRG), craniofacial derivatives, and neurectoderm derivatives (e.g., midbrain roof, thalamus) (Fig. 3f). To improve resolution, we fixed and sectioned several littermates and performed *in situ* hybridization for *ZsGreen,* identifying rare *ZsGreen* expression in the developing semilunar valves (Fig. 3g).

The labeling of neural crest structures with the *PthlhCreER* mouse line raised the interesting possibility that the *Pthlh*^+^ sub-population of endocardial cells may have a neural crest origin even though all endocardium has previously been thought to be derived from cardiac mesoderm. In agreement with this hypothesis, examination of the differentially expressed genes between *Pthlh*^+^/*Cdh5*^+^ and *Pthlh*^−^/*Cdh5*^+^ endocardium revealed an up-regulation of a number of neural crest-expressed genes, including *Stmn2*^29^, *Nts*^30^, and *Prnp*^31^, with UMAPs showing specific expression of these genes observed in *Pthlh*^+^ cells (Extended Data Fig. 4d). To test the hypothesis that *Pthlh^+^* cells are neural crest-derived, we reprocessed a previously published scRNA-seq dataset generated using FACS-sorted embryonic cells from the neural crest-specific Wnt1-Cre mouse line crossed with a tdTomato^+^ reporter mouse^32^. These data were generated using FACS-sorted tdTomato^+^ neural crest-derived cells from pooled, microdissected embryonic hearts at several developmental timepoints. We focused our analysis on the E17.5 timepoint, searching for cells with a similar transcriptional signature compared to the *Pthlh*^+^ cells we identified in our own scRNA-seq experiments. In E17.5 embryos, we identified a subset of tdTomato^+^ neural crest-derived cells positive for both *Pthlh* and the neural crest-enriched gene *Stmn2*, consistent with a neural crest origin for *Pthlh*^+^ cells (Extended data Fig. 4e). At E12.5, *Stmn2* transcripts were localized not only to the same valvular endocardial cells as *Pthlh*, but also to the neural tube, dorsal root ganglia, and peripheral nerves, representative of their neural crest origin (Fig. 3h).

### Neural Crest-Derived *Pthlh* Signaling is Required for Valvular Interstitial Cell Maturation

To more definitively evaluate the lineage origin of *Pthlh*^+^ endocardial cells, and the importance of *Pthlh* within the neural crest derivatives during valvulogenesis, we performed tissue-specific deletion of *Pthlh* using the *Wnt1*^Cre2^ mouse line^33^ crossed with a floxed allele of *Pthlh* (Fig. 4a). Similar to the whole organism *Pthlh* knockout, neural crest-specific *Pthlh* deletion resulted in perinatal cyanosis and lethality, with no *Wnt1*^Cre/+^:*Pthlh*^fl/fl^ (ΔNC-*Pthlh*) animals surviving to P7, whereas normal Mendelian ratios were observed at E18.5 (Extended Data Fig. 5a). To be certain neural crest-derived *Pthlh* was essential, we deleted *Pthlh* with a second neural crest-specific Cre using the *Sox10^Cre^* mouse^34^; again, we observed a loss of Mendelian ratios by P7 with no *Sox10^Cre^*:*Pthlh*^fl/fl^ surviving to weaning age (Extended Data Fig. 5b). Thus, neural crest-derived *Pthlh* signaling appears to be required for survival.

**Figure 4.**
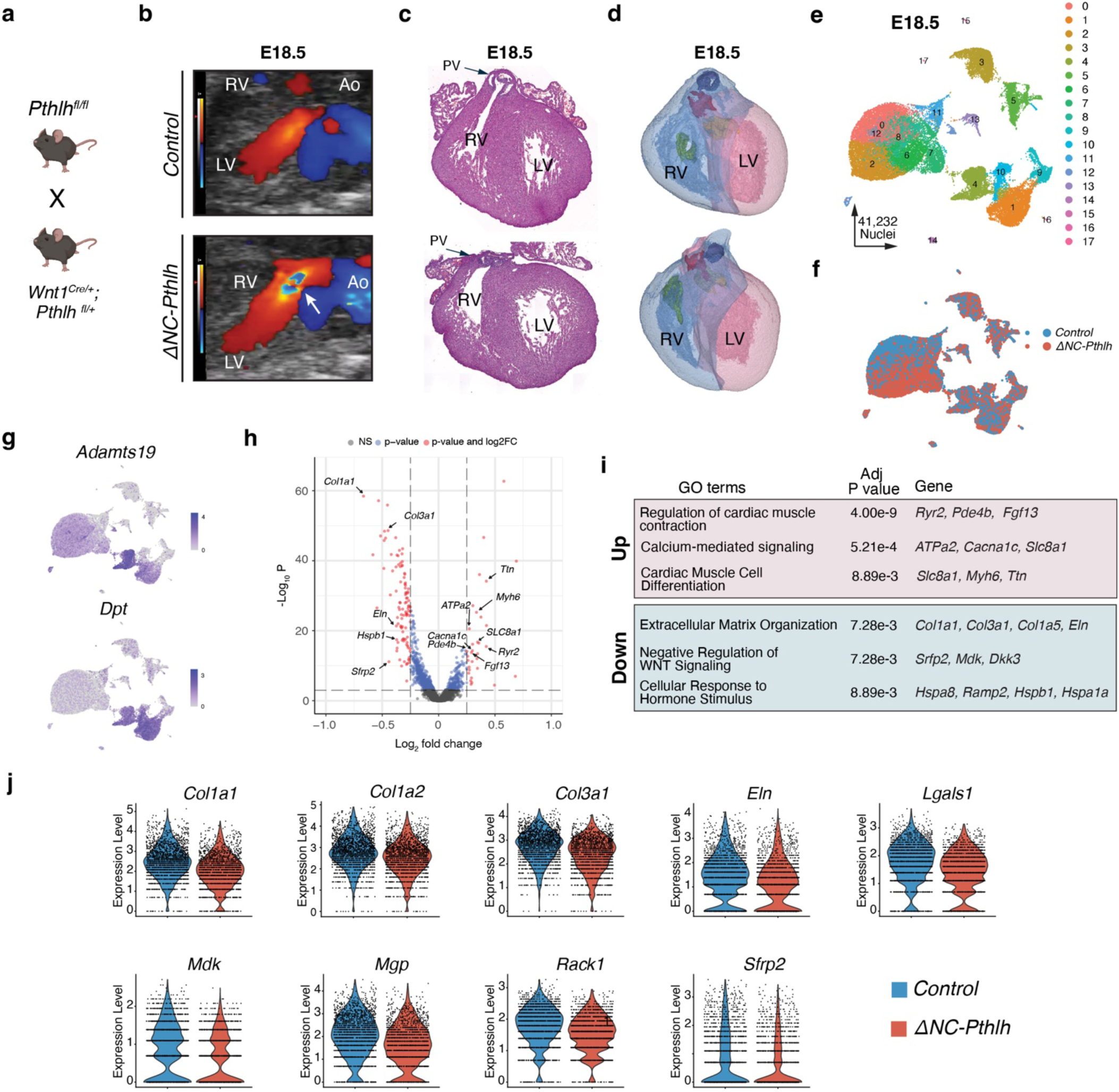
Neural crest-derived *Pthlh* signaling is essential for valvular interstitial cell maturation. **a**, Mating strategy for deletion of neural crest-derived *Pthlh* using *Pthlh^fl/fl^* females and Wnt1^Cre/+^; Pthlh^fl/+^ males. **b**, Fetal echocardiogram from control (top) and ΔNC-*Pthlh* (bottom) E18.5 hearts. **c**, Hematoxylin and eosin staining of control (top) and ΔNC-*Pthlh* (bottom) E18.5 hearts, with a dysplastic pulmonary valve (PV) shown (Black arrow). **d**, Three-dimensional reconstruction of control (top) and ΔNC-*Pthlh* (bottom) E18.5 hearts from microCT data. **e**, Annotated UMAP plot from snRNA-seq of whole mouse hearts at E18.5 (n = 6, 3 control and 3 ΔNC-*Pthlh*), colored by cluster identity. **f**, Annotated UMAP plot from snRNA-seq of whole mouse hearts at E18.5 (n = 6, 3 control and 3 ΔNC-*Pthlh*), colored by condition (control in blue, ΔNC-*Pthlh* in red). **g**, UMAP plot of valvular interstitial cell markers *Adamts19* and *Dpt* in cluster 4. **h**, Volcano plot of differentially expressed genes in valvular interstitial cells (cluster 4) between control and ΔNC-*Pthlh* conditions (adjusted *P* value < 0.05 & Log2FC > 0.25). **i**, Top GO terms enriched in up- and down-regulated genes with representative genes with corresponding adjusted *P* values for each GO term represented in adjacent boxes. **j**, Violin plots for selected differentially expressed genes between control (blue) and ΔNC-*Pthlh* (red) conditions.

Fetal echocardiography at E18.5 showed turbulent flow through the aortic and pulmonary valves in ΔNC-*Pthlh* embryos, indicative of congenital valvular stenosis (Fig. 4b). Histologic analysis of ΔNC- *Pthlh* E18.5 embryonic hearts revealed a valvular dysplasia phenotype similar to the *Pthlh*^+^ cell ablation phenotype, characterized by thickened, immature-appearing leaflets. The penetrance of pulmonary or aortic valve dysplasia was 87.5% or 62.5%, respectively (Fig. 4c, Extended Data Table 2). Moreover, three-dimensional reconstruction of microCT data from ΔNC-*Pthlh* and control littermates revealed a highly dysplastic pulmonary valve phenotype (Fig. 4d). Thus, loss of neural crest-derived *Pthlh* signaling phenocopies a significant portion of the phenotype observed in the *Pthlh*^+^ cell ablation model, providing evidence of neural crest *Pthlh*-mediated regulation of valve development.

To define the transcriptional changes associated with the valvular defects induced by loss of neural crest-derived *Pthlh* signaling, we performed single nuclear RNA sequencing (snRNAseq) on three E18.5 *Wnt1^Cre+/-^*:*Pthlh ^fl/fl^* (ΔNC-*Pthlh)* samples and three E18.5 *Wnt1^Cre-/-^;Pthlh ^fl/+^* (control) samples. A total of 41,232 high quality nuclei across 18 different clusters were analyzed and annotated using previously-established marker genes (Fig. 4e). We observed a substantial separation between the ΔNC-*Pthlh* and Control nuclei in the myocardial clusters (Fig. 4f), likely reflecting the myocardial remodeling secondary to valvular dysfunction. We focused our analysis primarily on the valvular interstitial cells (VICs), as these are putatively the principal target of the neural crest-derived *Pthlh* paracrine signal based on expression of its receptor, *Pth1r*, at this stage. Upon sub-setting and re-clustering of the mesenchymal population at E17.5, we identified six genes that had unique expression patterns within the E17.5 wild-type mesenchyme. *Adamts19*, *Dpt*, and *Cthrc1* reliably labeled specific regions within the developing valve mesenchyme, while *Ptprz1*, *Sfrp2*, and *Aspn* were not expressed within the valve mesenchyme (Extended Data Fig. 5c-g). Within the ΔNC-*Pthlh* animals, we observed an enrichment of *Adamts19^+^/Dpt^+^*cells in the VIC cluster (cluster 4) (Fig. 4g). Differential gene expression analysis within the VIC cluster expressing *Adamts19^+^/Dpt^+^* (cluster 4, 1797 ΔNC-*Pthlh* nuclei vs. 1908 control nuclei) identified a total of 149 genes significantly dysregulated within VICs upon loss of neural crest-derived endocardial *Pthlh* secretion (Fig. 4h). Down-regulated genes were associated with gene ontology terms, including extracellular matrix organization, negative regulation of WNT signaling, and cellular response to hormone stimulus (Fig. 4i). Three collagen isoforms (Col1a1, Col1a2, Col3a1) and elastin, which are known to be critical to VIC function, were downregulated, as well as an inhibitor of endochondral ossification, matrix Gla protein (*Mgp*) (Fig. 4j). Notably, Matrix Gla-Protein (*MGP)* expression is downregulated in calcified aortic valve interstitial cells in humans and deletion of this gene in mice leads to aortic valve calcification ^35,36^.

### Neural Crest-Derived *Pthlh* Signaling Acts as a BMP Modulator During Valve Morphogenesis

To dissect the mechanism by which VICs are dysregulated, we analyzed embryos at an earlier stage of development and performed snRNAseq on three ΔNC-*Pthlh* and two control hearts at E13.5, soon after *Pthlh* signaling becomes active during valvulogenesis (n=33,248 nuclei). We computationally subsetted the valvular cell populations (n=12,564), each labeled based on expression of the aforementioned marker genes, for further analysis (Fig. 5a-c, Extended Data Fig. 6a-d). Differential gene expression analysis between conditions revealed significant dysregulation of genes in valve interstitial and valve myocardial clusters (Fig. 5d,f; Extended Data Fig. 6e,f). Within valve myocardial cells, Bmp2 and Rspo3, myocardial-derived signals that promote EMT and cushion expansion, were upregulated in *Pthlh*-null cells. Correspondingly, gene ontology enrichment analysis showed downregulation of mesenchymal and cushion differentiation markers within *Pthlh*-null VICs, and upregulation of genes associated with skeletal development (Fig. 5e,g).

**Figure 5.**
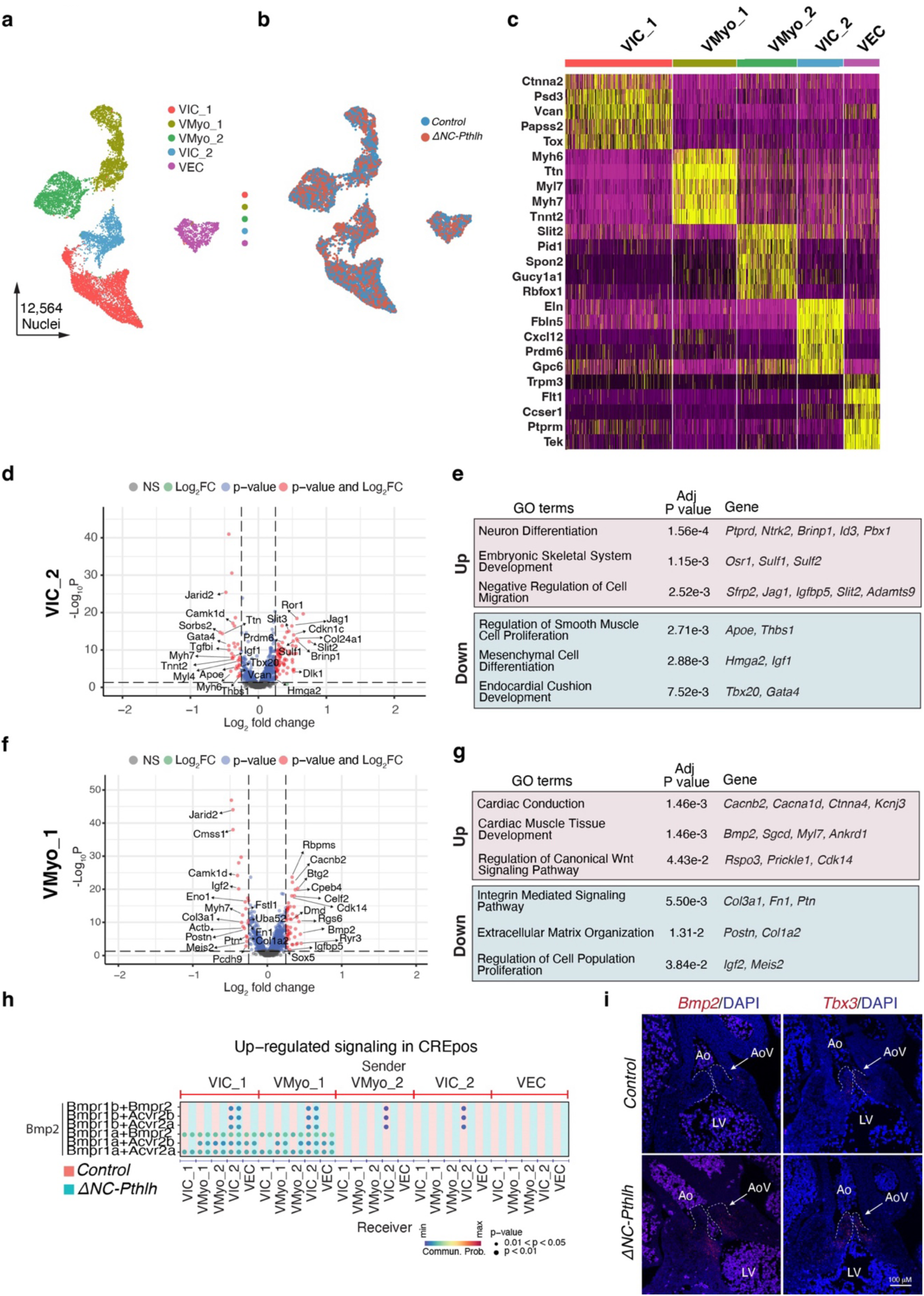
Neural crest-derived *Pthlh* signaling is a negative regulator of BMP signaling. **a**, Annotated UMAP plot of valvular cell types from snRNA-seq of whole mouse hearts at E13.5 (n = 5, 2 control and 3 ΔNC-*Pthlh*), colored by cluster identity. **b,** Annotated UMAP plot of valvular cell types from snRNA-seq of whole mouse hearts at E13.5 (n = 5, 2 control and 3 ΔNC-*Pthlh*), colored by condition (control in blue, ΔNC-*Pthlh* in red). **c**, Gene expression heatmap of marker genes for each identified valvular cell population at E13.5 (n = 5, 2 control and 3 ΔNC-*Pthlh*). **d,** Volcano plot of differentially expressed genes in valvular interstitial cell cluster 2 (VIC_2) between control and ΔNC-*Pthlh* conditions at E13.5 (adjusted *P* value < 0.05 & Log2FC > 0.25). **e,** Top GO terms enriched in up- and down-regulated genes in VIC_2 cells with representative genes with corresponding adjusted *P* values for each GO term represented in adjacent boxes. **f,** Volcano plot of differentially expressed genes in valvular myocardium cluster 1 (VMyo_1) between control and ΔNC-*Pthlh* conditions at E13.5 (adjusted *P* value < 0.05 & Log2FC > 0.25). **g**, Top GO terms enriched in up- and down-regulated genes in VMyo_1 cells with representative genes with corresponding adjusted *P* values for each GO term represented in adjacent boxes. **h,** Bubble plot of predicted up-regulated BMP ligand-receptor pairs between valvular cell types at E13.5 in the ΔNC-*Pthlh* condition. **i**, Fluorescent *in situ* hybridization for *Bmp2* (red) and *Tbx3* expression (red) in E13.5 outflow tracts in control (top row) and ΔNC-*Pthlh* (bottom row) embryos.

To identify paracrine signaling dysregulation secondary to the loss of neural crest-derived *Pthlh* signaling, we performed comparative cell-cell communication analysis between the ΔNC-*Pthlh* and control conditions via CellChat. Differential communication analysis revealed substantial changes in both the number and strength of interactions between the valvular cell types (Extended Data Fig. 6g). CellChat identified a number of pathways altered in a condition-specific manner, including the Notch, Neuregulin, and BMP signaling pathways (Extended Data Fig. 6h). Specifically, up-regulation of *Bmp2* – *Bmpr1a* signaling was suggested primarily from the valvular myocardium to the valvular interstitial cell populations (Fig. 5h). The major source of *Bmp2* during this stage of valve development is the valvular myocardium^37,38^, consistent with the upregulation of Bmp2 in the valvular myocardium upon *Pthlh* loss (Extended Data Fig. 6i). RNAscope correspondingly demonstrated an increase in *Bmp2* expression and its downstream valve myocardial target, *Tbx3,* at E13.5 in the setting of loss of neural crest-derived *Pthlh* (Fig. 5i).

Taken together, our work supports a model in which loss of *Pthlh* in neural crest-derived valve endocardium results in thickened, dysplastic valves with a concomitant increase in BMP expression and up-regulation of valvular myocardial gene programs, which may interfere with proper valvular interstitial maturation.

## Discussion

This study reveals a rare valvular subpopulation that is essential for valvular morphogenesis and implicates a novel peptide hormone signaling pathway in the context of heart development. The results in this study demonstrate a novel neural crest to endocardial transition, illustrating the significance of differential lineage contribution during valvulogenesis. Of note, recent clinical genetics studies have associated rare variants in multiple genes highly enriched in *Pthlh^+^*cells, including *PTPRB*^39^ and *WNT9B*^40^, with CHD phenotypes involving valve development. The genetic associations between these *Pthlh*^+^ cell marker genes and human CHD phenotypes emphasize the need to study this population of cells in the context of human development.

As it relates to *Pthlh* signaling, we initially found this pathway to be of interest due to the annotation of genes related to valve development, including *Msx2,* an important regulator of valvular mesenchyme differentiation^41,42^, and *Bmpr1a*^43^, loss of which in the developing atrioventricular myocardium results in a mouse model of Ebstein’s anomaly, a congenital valvulopathy defective in post-EMT valvulogenesis. Moreover, whole organism knockout of *Pthlh* results in perinatal cyanosis and lethality; the cause of this lethality was initially attributed to mechanical failure inhibiting respiration given the expression of *Pthlh* in the developing ribs^26^. However, in vivo echocardiography results presented here suggest that severe valve dysfunction also likely contributes to perinatal lethality. Our study reveals a novel molecular regulator of post-EMT valvulogenesis that acts to modulate BMP signaling, which plays a critical role in the proper maturation of valvular interstitial cells.

Autocrine BMP signaling is essential in the specification and maintenance of valvular myocardium through promotion of *Tbx2/Tbx3* expression, which drives valvular myocardial gene programs (e.g. *Rspo3)* and represses chamber-specific myocardial gene programs. In our perturbation model, loss of *Pthlh* resulted in increased valvular myocardial *Bmp2* expression and concomitant increase in downstream *Tbx3*. Our work supports a model in which *Pthlh* acts as a BMP repressive signal in the developing cushion, with loss of *Pthlh* resulting in increased BMP pathway activity, predicted Bmp2-Bmpr1a signaling, and increased *Tbx3* expression. Upregulation of this pathway correlates with impaired valvular interstitial maturation, and there may also be direct effects on VICs through Pth1r, ultimately contributing to immature, malfunctioning leaflets that remain thickened and stenotic.

These findings open several new investigational avenues related to valvular development and disease. The persistence of this developmentally specified cell type into adulthood raises important questions about its role in homeostasis and degenerative valve disease. Future studies of this rare cell type and Pthlh signaling will inform our understanding of congenital and adult valve disease pathogenesis, and improve our ability to diagnose, treat, and ultimately prevent valvular heart disease.

## Materials and Methods

### Animal Models

All mouse studies were carried out as stated our animal use protocol (AN189140) in accordance with all applicable ethical regulations including UCSF animal use guidelines, and the NIH Guide for the Care and Use of Laboratory Animals. The Institutional Animal Care and Use Committee granted approval for our animal use protocol. C57BL/6J wild type (Stock 000664), B6.Cg-Gt(ROSA)26Sortm6(CAG-ZsGreen1)Hze/J (Ai6) (Stock 007906), B6.Cg-Tg(Pthlh-cre/ERT2)909Nono/J (Pthlh^CreER^) (Stock 032873), B6.Cg-E2f1^Tg(Wnt1-cre)2Sor^/J (Wnt1^Cre^) (Stock 022501) mice were purchased from Jackson Laboratory (Bar Harbor, ME), Pthlh flox allele mice^1^ were provided by Andrew Karaplis’ lab.

### Embryo Dissection for RNAscope and Spatial Transcriptomics Assay

Embryos used for these assays were collected through timed mating similar to methods for single cell processing. Dissected tissues were fixed overnight with 4% paraformaldehyde (50-980-487, Fisher Scientific). Tissues were subsequently washed in PBS and dehydrated in 70% EtOH overnight. Tissues were processed in paraffin overnight and sectioned onto glass slides at 5 υm thickness.

### RNAscope In Situ Hybridization

Section RNA in situ hybridization was performed in using Multiplex Fluorescent Reagent Kit v2 (ACD, 323100) as previously reported. Each experiment was repeated using at least three biological replicates. The *in situ* hybridization probes used in this study are as follows: Pthlh (ACD, 456521), Dhh (ACD, 415031), Wnt9b (ACD, 405091), Aspn (ACD, 502051), Ptprz1 (ACD, 460991), Cthrc1 (ACD 413341), Dpt (ACD, 561511), Sfrp2 (ACD, 400381), Adamts19 (ACD, 500981), ZsGreen (ACD 461251), and Stmn2 (ACD, 498391), Bmp2 (ACD, 406661), Tbx3 (ACD, 438211). Spatial transcriptomics was performed in FFPE sections using the Xenium Prime 5K Mouse Pan Tissue & Pathways Panel and analyzed with Xenium Analyzer, following manufactures’ instruction (10XGenomics). All slides were imaged on Olympus FV3000RS confocal microscope.

### Light sheet Microscopy

For light sheet microscopy imaging, the whole mount samples were microdissected to keep the area of interest. Samples were fixed in 1% paraformaldehyde for 15 minutes at room temperature. Samples were then washed three times in PBS. Samples were incubated overnight at 37°C in CUBIC-L solution (TCI chemicals, T3740). After validating that the sample had been cleared, samples were washed three times for one hour in PBS at 37°C. Samples were then incubated overnight in 50% PBS/ 50% Cubic R+ (M) solution (TCI chemicals, T3741). Samples were subsequently placed in 100% Cubic R+ (M) solution in a dark area until imaging. Samples were then imaged on a SmartSPIM Light Sheet Microscope (LifeCanvas Technologies, Cambridge). Images were processed with Fiji and Imaris (Oxford Instruments).

### Fetal echocardiography by trans-uterine doppler analysis

Timed-pregnant mice at embryonic day 18.5 (E18.5) were anesthetized with 1–2% isoflurane in oxygen and positioned supine on a temperature-controlled platform (37 °C). Maternal heart rate, respiratory rate, and temperature were continuously monitored throughout the procedure. To maintain hydration and uteroplacental perfusion, pre-warmed sterile saline was applied intermittently to the uterus during imaging. Following aseptic preparation of the maternal abdomen, a midline laparotomy was performed to gently exteriorize the uterine horns. The uterus was carefully positioned on a pre-warmed saline-soaked gauze pad to preserve temperature and moisture, while keeping the umbilical circulation and maternal blood supply intact. Only a limited segment of the uterus was exposed at any time to minimize heat and fluid loss. Embryos were imaged in situ through the intact uterine wall and amniotic sac. Pre-warmed ultrasound gel was applied to maintain acoustic coupling and prevent hypothermia. Fetal echocardiography was performed using a high-frequency ultrasound imaging system (Vevo 3100, FUJIFILM VisualSonics) equipped with a 30–40 MHz linear array transducer. Image acquisition was conducted in B-mode, M-mode, and pulsed-wave (PW) Doppler modes. Individual embryos within the uterus were visualized in B-mode by scanning transversely through the maternal abdomen. Embryos were selected based on accessibility and stable heart rate. The imaging probe was gently adjusted to obtain an optimal four-chamber or long-axis view of the fetal heart. Pulsed-wave Doppler recordings were obtained to detect the presence of AV shunts. The Doppler sample volume (≈0.1– 0.2 mm) was positioned parallel to the direction of flow to minimize angle error (<15°). After imaging, the mother was sacrificed and embryos collected for micro-CT analysis and histology as described.

### Micro-computed tomography (micro-CT)

E18.5 embryonic hearts were collected and fixed in 4% paraformaldehyde overnight at 4 °C. Samples were rinsed in cold 1X phosphate-buffered saline (PBS) twice followed by dehydration in cold 70% ethanol (EtOH). Store the samples at 4 °C until further processing. Incubate the samples in 1% phosphotungstic acid (PTA, Sigma P4006-10G) in 70% EtOH solution at 4 °C for at least 72 hours. Samples were then washed with 70% EtOH to remove excess PTA and then were embedded in 0.5% agarose gel (UltraPureTM Agarose, Invitrogen, Cat No. 15510-027) in 15ml Falcon tubes. Samples were kept on ice during transportation and pursue with scanning at the UCSF Skeletal Biology & Biomechanics Core facility (μCT 50 cabinet microCT scanner, SCANCO Medical AG. Switzerland). The images were processed with OsiriX MD (Pixmeo SARL). Collected hearts were also collected for histology, followed by H&E staining for heart analysis. All conclusions for presence of septal and valvular defects were performed double-blinded by two researchers independently.

### 3-Dimensional Heart Model Generation

To create the cardiac 3D models, we loaded the micro-CT dataset into Materialise Mimics Medical version 25.0. Cardiac regions of interest (ROI), ex: atria, ventricles, valves, inflows/outflows, arteries and veins, etc were labeled through a process of segmentation and multiple 3D objects were generated corresponding to these ROIs. CT-based Hounsfield Units (HU) threshold windows were selected to include the relevant anatomy. The resulting voxel (3D pixel) masks were further post-processed in Materialise 3-Matic version 17.0 and converted to surface-rendered 3D mesh models. Thereafter a comprehensive digital 3D model was finalized.

### Human fetal heart spatial transcriptomics

Visium HD data from the human developing heart atlas^22^ were obtained from the Heart Cell Atlas (heartcellatlas.org). We used the authors’ processed objects, in which 2-um bins are aggregated into cells with bin2cell ^44^ and tissue niches are defined with CellCharter^45^, as described in the source publication. Analysis was performed on the 9 PCW section (donor C194; 136,685 cells). Cells with >= 2 UMI for PTHLH or CDH5 were plotted on the section coordinates with the authors’ niche annotation. Analysis and plotting were performed in R 4.5 (ggplot2, data.table, scattermore).

### Fetal valve cell isolation

Fetal heart samples were obtained directly from the operating room and placed in 50 mL conical tubes containing PBS (Life Technologies, 14190250) with 1% FBS (ThermoFisher Scientific, 10439016) and heparin (100 µL heparin per 50 mL PBS/FBS), and transported to the dissection room on ice. Samples were dissected under a dissection scope, and regions of interest were isolated using micro tweezers, micro scissors, and a micro scalpel, and transferred into 1.5 mL Eppendorf tubes containing PBS with 1% FBS; remaining tissue was flash frozen and stored at −80°C for future experiments. Samples from each valve and each chamber were individually isolated. Dissected cardiac tissue was mechanically torn into several pieces with micro scissors prior to incubation at 37°C in 300 µl 1 mg/mL Liberase TL (Roche, 05401020001) for 5 min, dissociated for an additional 10 min with intermittent trituration, and quenched with 900 µl PBS with 1% FBS. Cells were filtered through a FACS tube filter, centrifuged at 150g for 3 min, and resuspended in PBS with 1% FBS. Cell concentration was determined manually by hemocytometer using trypan blue exclusion. For scRNAseq, the cell suspension was prepared to recover 10K cells per sample according to the manufacturer’s instructions (Chromium Single Cell 3’ Reagent Kit v3.1, 10X Genomics).

### Single-cell Isolation, Library Generation, and Sequencing

For all timed mating experiments, male and female mice were housed overnight and female mice were visually inspected for a vaginal plug in the morning. Embryos at date of plug were considered embryonic day 0.5 (E0.5). Embryos from E12.5, E13.5, and E17.5 were prepared in the same manner. Embryos were dissected in cold PBS (Life Technologies, 14190250) with 1%FBS (ThermoFisher Scientific, 10439016). Dissected cardiac tissue was mechanically torn into several pieces with forceps prior to incubated at 37°C in 200 µl 0.25 mg/mL Liberase (Roche, 05401020001) for 5 min, dissociated for 15 min, and quenched with 600 µl PBS with 1% FBS. Cells were filtered through a 70-µm cell strainer (BD Falcon, 08-771-2), centrifuged at 150g for 3 min, and resuspended in PBS with 1% FBS. For scRNAseq, the cell suspension was prepared to recover 10K cells per sample according to the manufacturer’s instructions (Chromium Single Cell 3’ Reagent Kit v.3, 10X Genomics). For the 10X Genomics Gene Expression and Chromatin Accessibility Co-Assay (scMultiome), the single-cell suspension was subjected to nuclear isolation according to the 10x nuclei isolation protocol (Nuclei Isolation for Single Cell ATAC Sequencing, CG000169, 10X Genomics). GEM generation for both assays was performed on the Chromium controller, using Chip G (PN-1000120) for scRNAseq and Chip J (PN-1000234) for scMultiome. The scRNAseq and scMultiome library preparations were performed according to the protocols described above using Chromium Single Cell 3ʹ Reagent Kits v3.1 (PN-1000121) and Chromium Next GEM Single Cell Multiome ATAC + Gene Expression Reagent Bundle (PN-1000283), respectively. All libraries were sequenced on the NovaSeq S4 200, in accordance with the manufacturer’s protocol.

For the Wnt1^Cre^;Pthlh^fl/fl^ Multiome experiment, embryo tails were dissected placed in QuickExtract DNA Extraction Solution (Lucigen, QE09050) and processed according to manufacturer’s protocols. Genotyping was performed by PCR using Phire Green Hot Start II DNA Polymerase (Thermo Fisher Scientific, F124L), according to manufacturer’s protocols, using JAX protocol primers for Wnt1^Cre^ and the following for Pthlh^fl^ allele (P1: GAGGCTAAGCCAGGAGGATT; P2: CCCCATCCTCTCTCCTCTCT). The band size for the wild type gene is 150bp and the band size for the floxed gene is ∼270bp.

### Single-cell Seurat Object Preparation

Raw sequencing data were preprocessed with the Cell Ranger v.5.0.0 pipeline for scRNAseq or Cell Ranger Arc for scMultiome (10X Genomics). Data from all samples, per experiment, was merged via cellranger-aggr or cellrangerarc-aggr and normalized to the same sequencing depth, resulting in a single gene-barcode matrix. Further analysis was performed using Seurat v4.0 R package. Quality control metrics used to remove low quality cells were: number of genes per cells between 4,000 and 9,500, UMI count per cell between 10,000 and 100,000, and mitochondrial gene percentage between 1 and 6%. Cell cycle scores were calculated and added to the metadata for each cell and the SCTransform function was used for normalization against cell cycle score. Clustering of cells was performed with the following commands: RunPCA, RunUMAP, FindNeighbors, and FindClusters. To identify marker genes, the clusters were compared pairwise for differential gene expression using the FindAllMarkers function (min.pct = 0.45, logfc.threshold=0.4, and return.thresh (p-value cut-off) = 1 × 10^−10^). Annotation of clusters was achieved via cross-referencing the resulting cluster-specific marker genes with known cell type-specific maker genes in the literature, we classified them into clusters of seven cell types. For visualization of gene expression data, the following Seurat functions were used: FeaturePlot, VlnPlot, DotPlot, and DoHeatmap.

### Cell-cell communication analysis

Analysis was performed using the R package CellChat. Seurat objects containing specific cell types and time points of interest were imported to CellChat using the function GetAssayData and createCellChat. All ligand-receptor and signaling pathways within the CellChatDB.mouse were kept for analysis. Initial pre-processing to identify over-expressed ligands and receptors was performed using the functions identifyOverExpressedGenes and identifyOverExpressedInteractions with the same settings as stated in the tutorial. Inference of cell communication was calculated with computeCommunProb and filtered by filterCommunication. Pathway level cell communication was calculated with computeCommunProbPathway and aggregated networks were identified with aggregateNet, using standard settings. Network centrality scores were assigned with the function netAnalysis_computeCentrality. This workflow was run for both control and neural crest-specific *Pthlh*-knockout datasets independently and differential signaling analysis was then run by merging the objects with mergeCellChat. A heatmap of differential signaling, based on significantly identified pathways from above, was generated using the function netAnalysis_signalingRole_heatmap and restricting the visualization to selected pathways.

## Acknowledgements

We are grateful to the family who consented to donate tissue for this study, without whom this research would not have been possible. We thank Peter Kouretas, MD, PhD, Shunji Sano, MD, PhD, Andrew Connolly, MD, PhD, Anita Moon-Grady, MD, and Jeffrey Bush, PhD, for their thoughtful discussion and critical insight throughout this project. We thank Tami Tolpa for her assistance with scientific illustrations and Bethany Taylor for editorial assistance. We thank the Gladstone Histology and Light Microscopy Core, the UCSF Center for Advanced Technology (CAT), and the Skeletal Biology and Biomechanics Core for their assistance with this work.

## Author contributions

A.F.M. and D.S. designed and directed all experiments in this study. A.F.M., M.W.C., S.S.R., and C.A.G. contributed to experimental design and analysis of single-cell experiments. A.F.M., A.P., and C.J.Y. performed computational analysis of single-cell experiments. A.F.M., M.W.C., T.N., and F.L. performed lightsheet microscopy, with A.F.M., M.W.C., and T.N. also performing histologic analysis. Y.H. performed fetal echocardiography. A.F.M. and A.L. performed mouse breeding and husbandry. A.F.M., A.B., and V.Y. performed RNAscope in situ hybridization. F.G. performed spatial transcriptomics. A.F.M. performed micro-CT, and S.A. and M.B. performed three-dimensional reconstruction analysis. A.F.M., I.K., and B.B. contributed to human fetal heart tissue collection, processing, and data analysis. D.S. and C.J.Y. supervised the project. A.F.M., M.W.C., and D.S. wrote the manuscript with input from all authors.

---

**Extended Data Figure 1.**
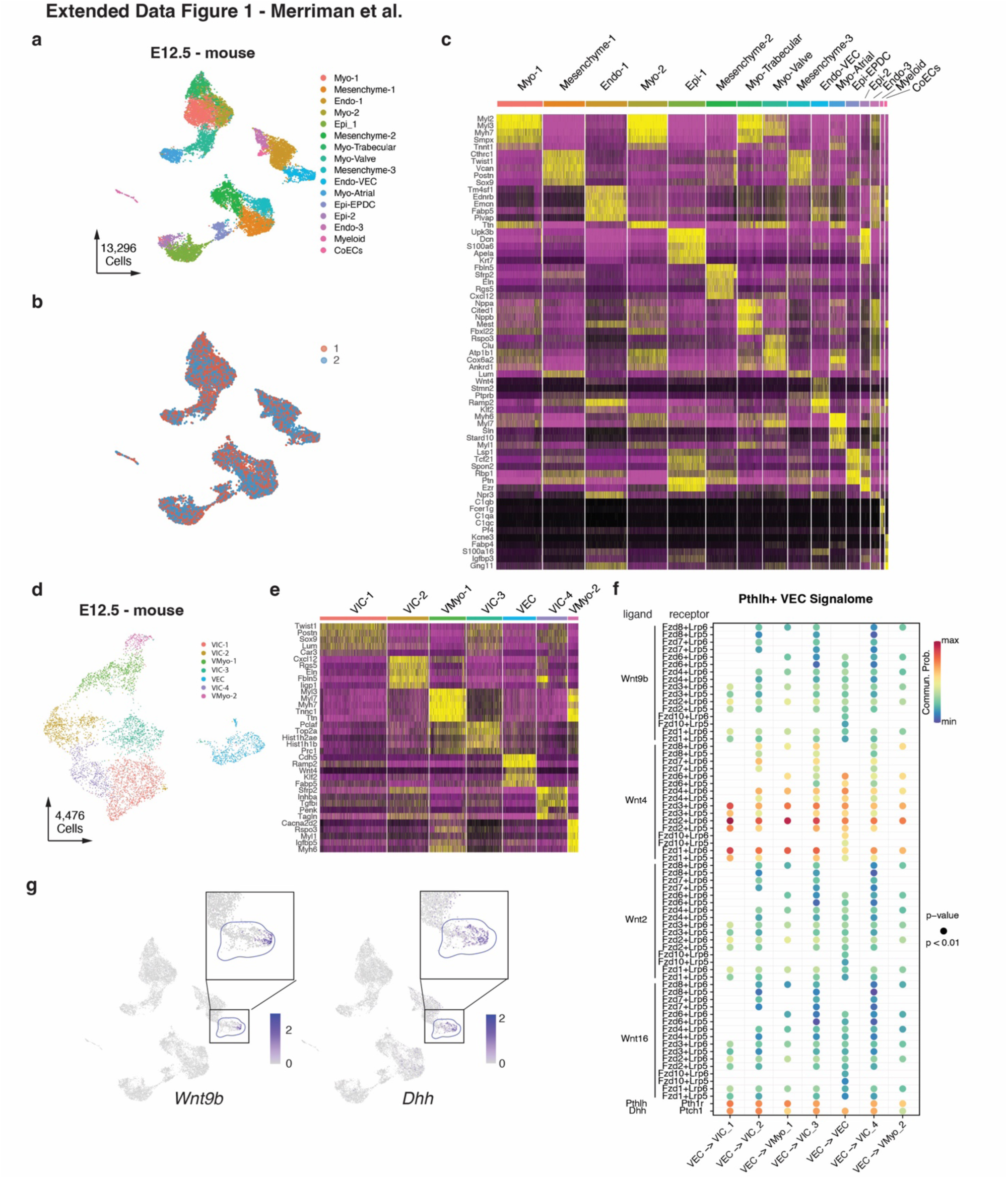
**a**, Annotated UMAP plot from scRNA-seq of whole mouse hearts at E12.5 (n = 2 WT), colored by cluster with corresponding cluster identities. **b**, UMAP plot of whole mouse hearts at E12.5 (n = 2 WT), colored by sample identity. **c**, Gene expression heatmap of marker genes for each identified cell population in the whole mouse heart at E12.5. **d,** Annotated UMAP plot of re-clustered valvular cell types at E12.5, colored by cluster with corresponding cluster identities. **e**, Gene expression heatmap of marker genes for each identified valvular cell population at E12.5. **f,** Bubble plot of predicted ligand-receptor pair interactions for the WNT, HH, and *PTHLH* signaling pathways between the VEC (sender) and valve mesenchyme and myocardium (receivers). **g**, UMAP plots of *Wnt9b* and *Dhh* expression with the valvular endocardial cell cluster highlighted (blue dashed circles) and zoomed in views in the right upper corners.

**Extended Data Figure 2.**
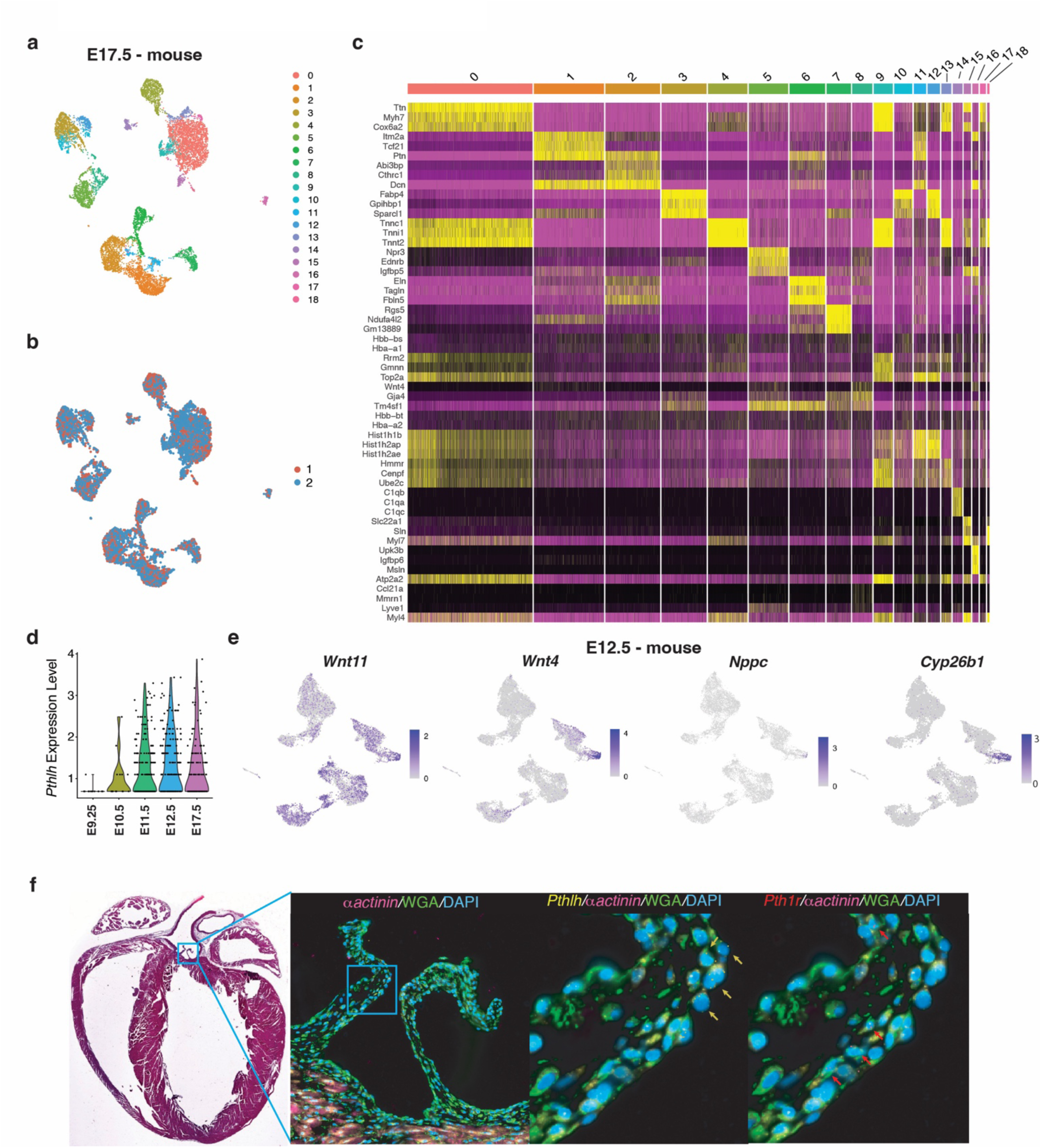
**a**, Annotated UMAP plot from scRNA-seq of whole mouse hearts at E17.5 (n = 2 WT), colored by cluster with corresponding cluster identities. **b**, UMAP plot of whole mouse hearts at E17.5 (n = 2 WT), colored by sample identity. **c**, Gene expression heatmap of marker genes for each identified cell population in the whole mouse heart at E17.5. **d**, Violin plot of *Pthlh* expression split by timepoint. E9.25 data were reprocessed from De Soysa et al., E10.5 & E11.5 were reprocessed from Ranade et al. **e**, UMAP plots of *Wnt11, Wnt4, Nppc,* and *Cyp26b1* expression in developing whole mouse heart at E12.5. **f**, Spatial transcriptomics of mouse adult hearts showing highlighting *Pthlh* (yellow dots and arrows) and Pth1r (red dots and arrows) expression in aortic valves. Cardiomyocytes are labelled by a-actinin (magenta), cellular membrane by WGA (green) and nucleus by DAPI (blue).

**Extended Data Figure 3.**
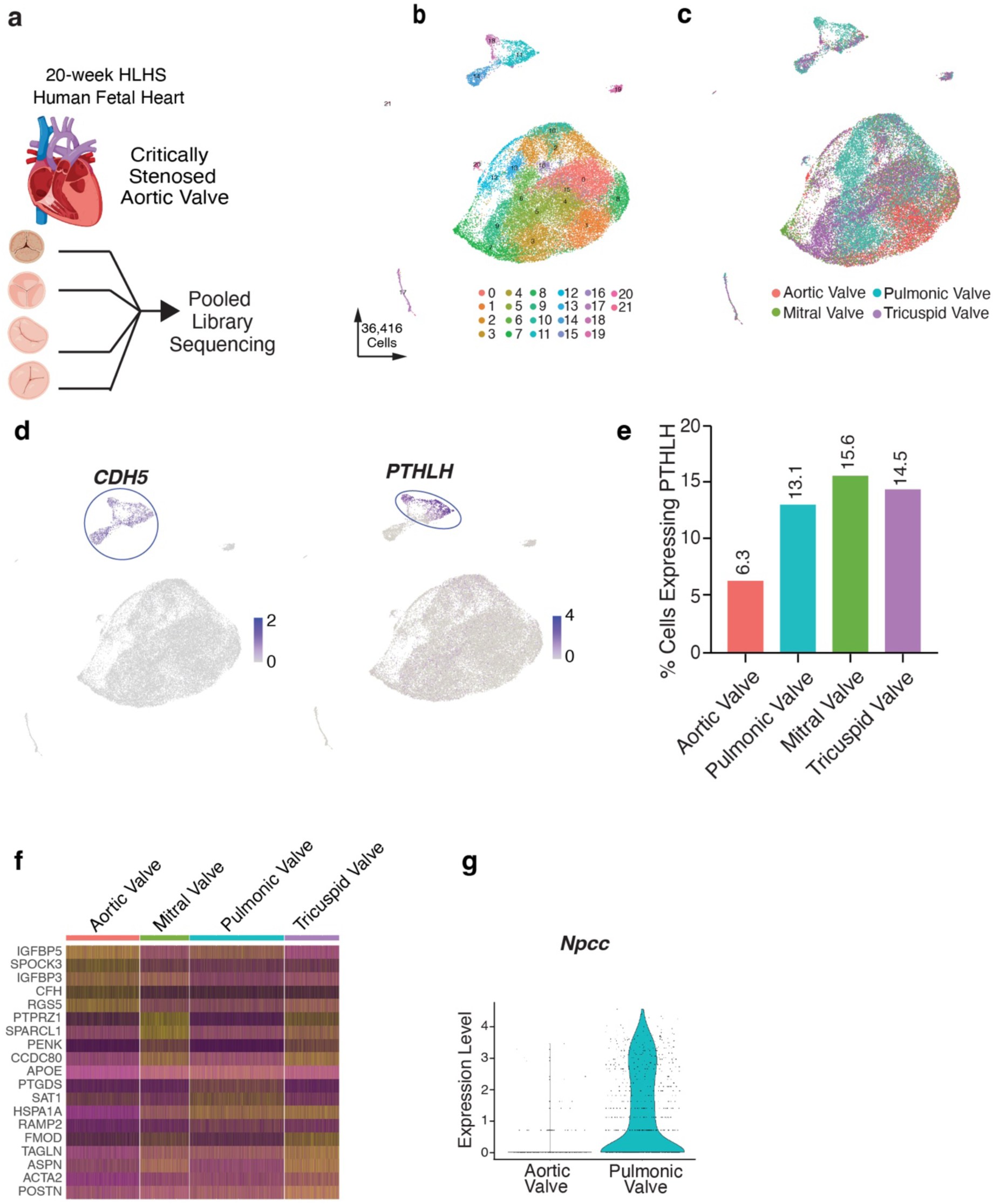
**a**, Cartoon representation of a scRNAseq experiment performed on each of the four cardiac valves from a 20-week human fetal heart with hypoplastic left heart syndrome and critical aortic stenosis. **b,** UMAP plot of all captured valvular cell populations from a 20-week human fetal heart with hypoplastic left heart syndrome and critical aortic stenosis colored by cluster. **c,** UMAP plot of all captured valvular cell populations colored by cardiac valve sample identity. **d,** UMAP plot of *CDH5* and *PTHLH* expression from a 20-week human fetal heart with hypoplastic left heart syndrome and critical aortic stenosis., **e,** Bar plot representing the percentage of cells in each valve expressing *PTHLH*. **f,** Gene expression heatmap plot of top 5 marker genes for each valve. **g,** Violin plot of NPPC expression between diseased aortic valve versus pulmonic valve cells.

**Extended Data Figure 4.**
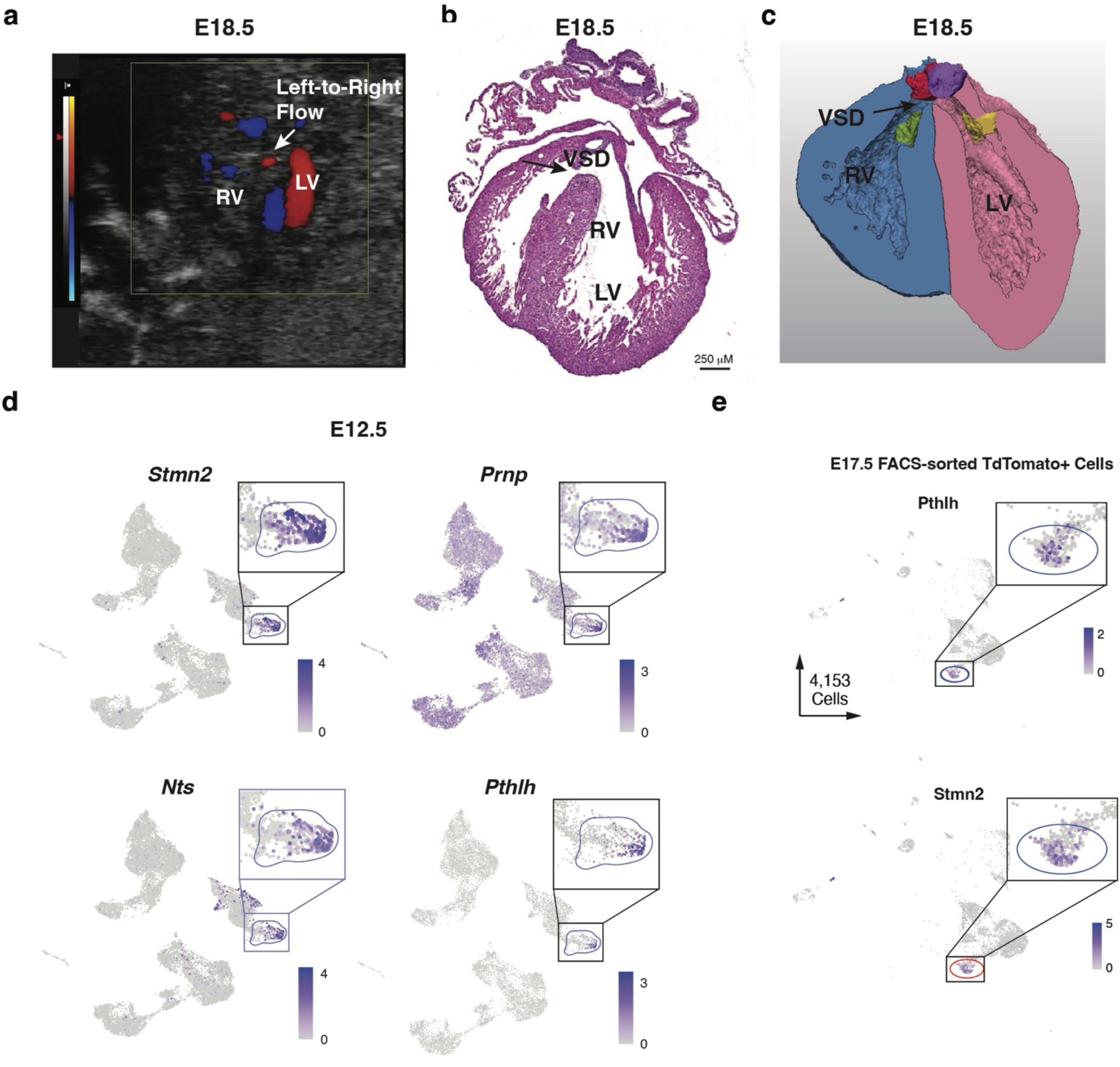
**a**, Fetal echocardiogram demonstrating left-to-right flow across a sub-aortic ventricular septal defect (VSD) in an E18.5 PthlhCreER/^+^; DTA/DTA embryo. **b,** Hematoxylin and eosin staining of an E18.5 PthlhCreER^+^ ablated heart demonstrating a sub-aortic VSD. **c**, Three-dimensional reconstruction of an E18.5 PthlhCreER^+^ ablated heart demonstrating a sub-aortic VSD from microCT data. **d,** UMAP plots of *Stmn2*, *Prnp*, *Nts*, and *Pthlh* expression in E12.5 wild-type whole mouse hearts with the valvular endocardial cell cluster highlighted (blue circle) and a zoomed in view in the right upper corner. **e**, *Pthlh, Stmn2* dual positive cells identified in re-processed scRNA-seq data from FACS-sorted neural crest-derived cells during cardiogenesis at E17.5 (Adapted from Chen et al, 2021).

**Extended Data Figure 5.**
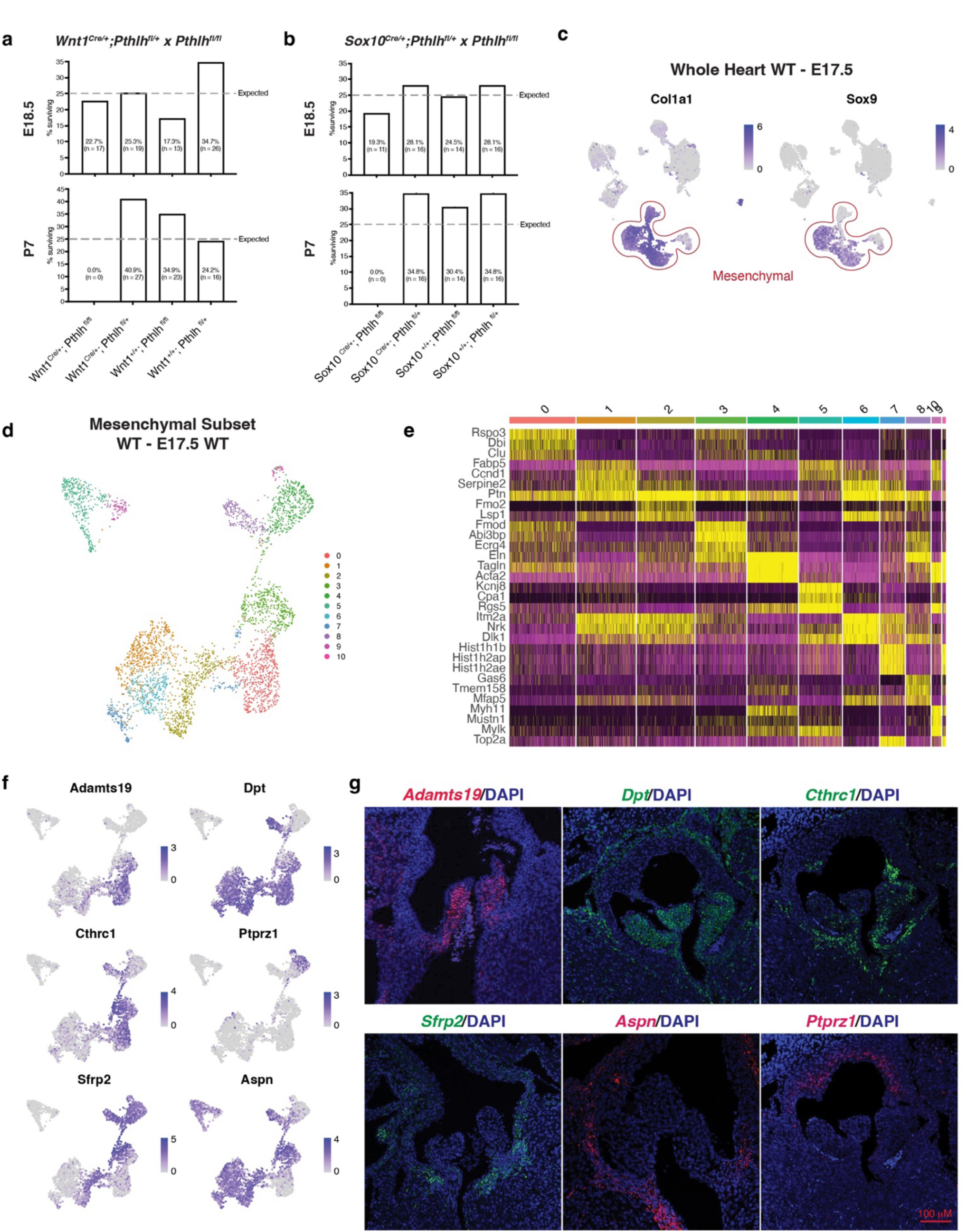
**a**, Chart demonstrating observed genotype frequencies for the Wnt1^Cre/+^;Pthlh^fl/+^ x Pthlh^fl/fl^ mating at E18.5 (top) and P7 (bottom). b, Chart demonstrating observed genotype frequencies for the Sox10^Cre/+^;Pthlh^fl/+^ x Pthlh^fl/fl^ mating at E18.5 (top) and P7 (bottom). **c**, UMAP plots of *Col1a1* and *Sox9* expression in E17.5 wild-type whole mouse hearts marking mesenchymal cell types. **d**, Annotated UMAP plot of computational subset and re-clustered mesenchymal cell types in E17.5 wild type whole mouse hearts, colored by cluster. **e**, Gene expression heatmap of marker genes for each identified mesenchymal cell population at E17.5. **f**, UMAP plots of *Adamts19, Dpt, Cthrc1, Ptprz1, Sfrp2,* and *Aspn* gene expression in E17.5 wild-type mouse mesenchyme. **g**, Fluorescent *in situ* hybridization for expression of (top row) *Adamts19* (red), *Dpt* (green), *Cthrc1* (green), and (bottom row) *Sfrp2* (green), *Aspn* (red), and *Ptprz1* (red) in the E17.5 wild-type aortic valve and para-valvular structures.

**Extended Data Figure 6.**
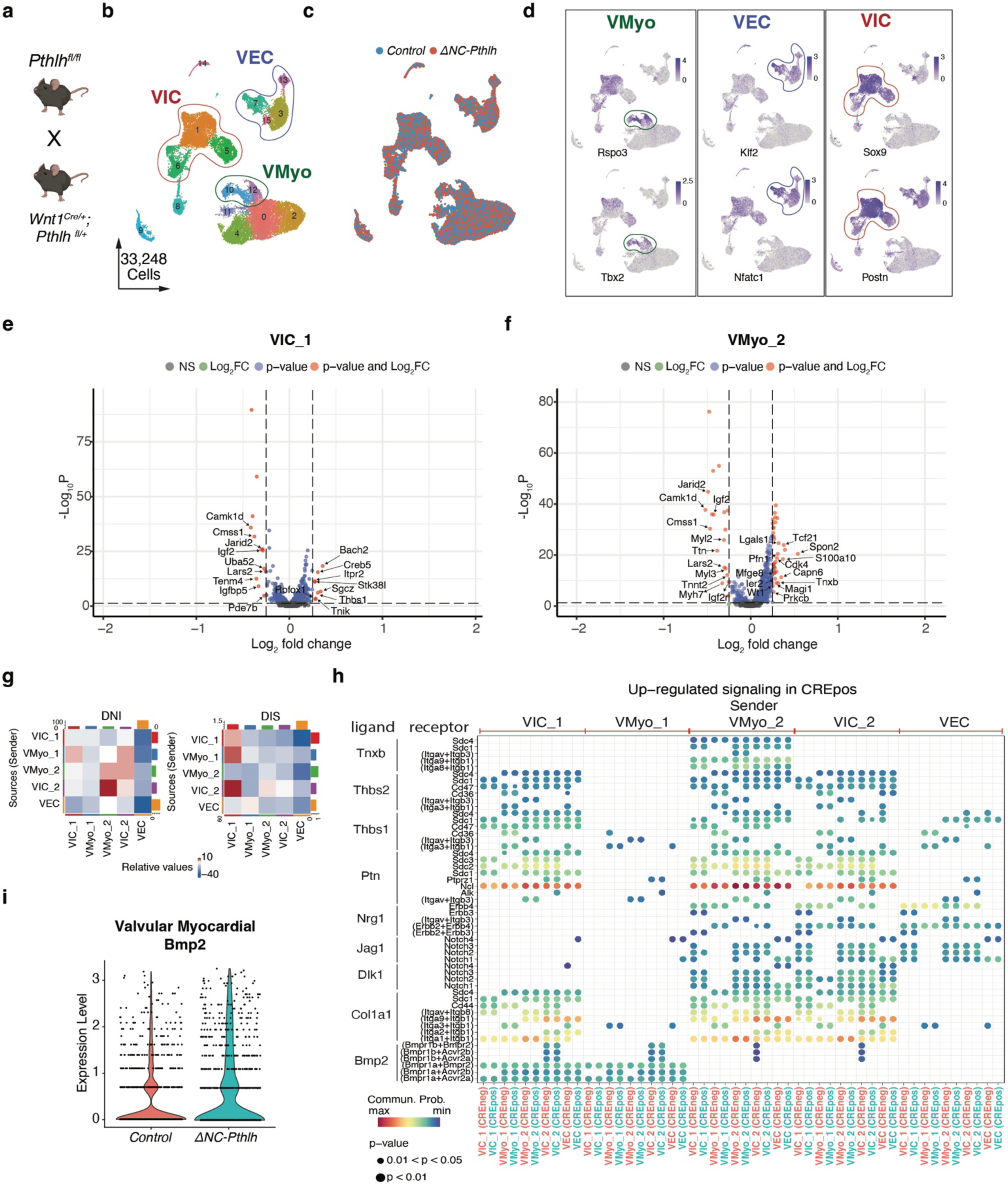
**a**, Mating strategy for deletion of neural crest-derived *Pthlh* using *Pthlh^fl/fl^* females and Wnt1^Cre/+^; Pthlh^fl/+^ males. **b,** Annotated UMAP plot of valvular cell types from snRNA-seq of whole mouse hearts at E13.5 (n = 5; 2 control and 3 ΔNC-*Pthlh*), colored by cluster with corresponding cluster identities, with the valvular myocardium (VMyo, green circle), valvular endocardial cells (VEC, blue circle), and valvular interstitial cells (VIC, red circle) highlighted. **c**, Annotated UMAP plot from snRNA-seq of whole mouse hearts at E13.5 (n = 5, 2 control and 3 ΔNC-*Pthlh*), colored by condition (control in blue, ΔNC-*Pthlh* in red). **d,** UMAP plots for previously characterized marker gene expression for *Rspo3, Tbx2, Klf2, Nfatc1, Sox9,* and *Postn*. **e,** Volcano plot of differentially expressed genes in valvular interstitial cell cluster 1 (VIC_1) between control and ΔNC-*Pthlh* conditions at E13.5 (adjusted *P* value < 0.05 & Log2FC > 0.25). **f,** Volcano plot of differentially expressed genes in valvular myocardium cluster 2 (VMyo_2) between control and ΔNC-*Pthlh* conditions at E13.5 (adjusted *P* value < 0.05 & Log2FC > 0.25). **g**, Heatmap plots showing differential number of interactions (DNI) and differential interaction strength (DIS) between valvular cell populations at E13.5 between control and ΔNC-*Pthlh* conditions. **h,** Bubble plot of predicted up-regulated ligand-receptor interactions between valvular cell types at E13.5 in the ΔNC-*Pthlh* condition. **i**, Violin plot of valvular myocardial *Bmp2* expression between control (red) and ΔNC-*Pthlh* (teal) conditions at E13.5.

**Extended Data Table 1.**

|  | **CONTROL (N=11)** | **ABLATION (N=15)** |
|---|---|---|
| **PULMONARY VALVE DYSPLASIA** | 1 (9.1%) | 12 (80.0%) |
| **AORTIC VALVE DYSPLASIA** | 1 (9.1%) | 11 (73.3%) |
| **MITRAL VALVE DYSPLASIA** | 2 (18.2%) | 4 (26.7%) |
| **TRICUSPID VALVE DYSPLASIA** | 1 (9.1%) | 3 (20.0%) |
| **VSD** | 1 (9.1%) | 4 (26.7%) |

**Extended Data Table 2.**

|  | **CONTROL (N=5)** | **ΔNC *Pthlh* (N=8)** |
|---|---|---|
| **PULMONARY VALVE DYSPLASIA** | 0 (0%) | 7 (87.5%) |
| **AORTIC VALVE DYSPLASIA** | 0 (0%) | 5 (62.5%) |
| **MITRAL VALVE DYSPLASIA** | 0 (0%) | 1 (12.5%) |
| **TRICUSPID VALVE DYSPLASIA** | 0 (0%) | 2 (25.0%) |
| **VSD** | 0 (0%) | 0 (0%) |

